# STARDUST: a pipeline for the unbiased analysis of astrocyte regional calcium dynamics

**DOI:** 10.1101/2024.04.04.588196

**Authors:** Yifan Wu, Yanchao Dai, Katheryn B. Lefton, Timothy E. Holy, Thomas Papouin

**Affiliations:** Washington University in St. Louis, Department of Neuroscience, St. Louis, MO 63110 USA

## Abstract

Calcium imaging has become a popular way to probe astrocyte activity, but few analysis methods holistically capture discrete calcium signals that occur across the astrocyte domain. Here, we introduce STARDUST, a pipeline for the Spatio-Temporal Analysis of Regional Dynamics & Unbiased Sorting of Transients from fluorescence recordings of astrocytes, and provide step-by-step guidelines. STARDUST yields fluorescence time-series from data-defined regions of activity and performs systematic signal detection and feature extraction, enabling the in-depth and unbiased study of astrocyte calcium signals.

**Graphical abstract:** 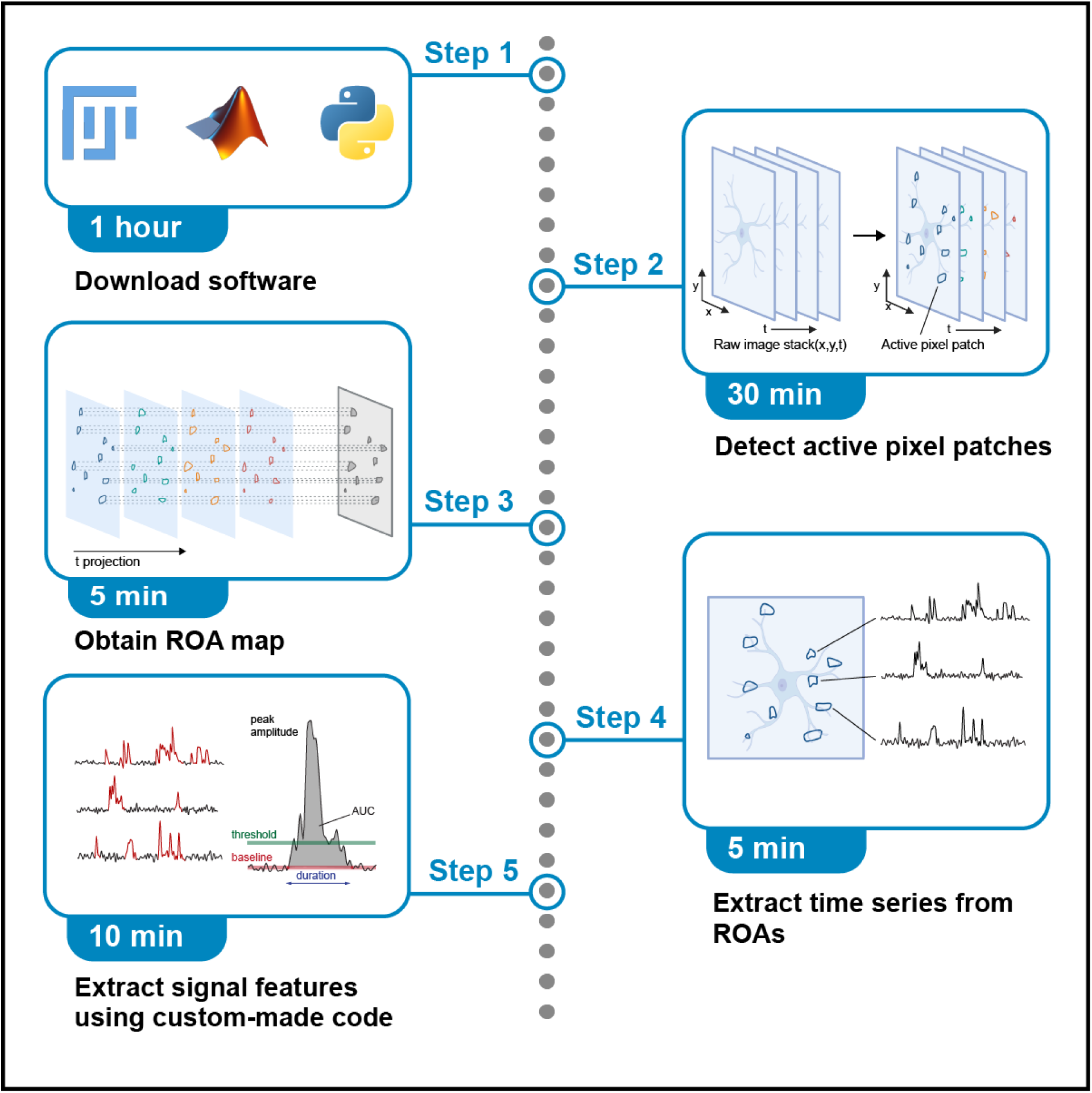

## Before you begin

Astrocytes, a main type of non-neuronal cells, display fluctuations of intracellular calcium concentrations ([Ca^2+^]i) that are responsive to ambient conditions and neuroactive molecules.^1–3^ Fluorescence imaging of Ca^2+^ indicators is thus a popular means to probe astrocyte activity in vitro, ex vivo and in vivo.^1–3^ Proven methods of neuronal Ca^2+^ imaging and analysis have been widely adapted to astrocytes for this purpose. However, important differences between neurons and astrocytes mean that key operational assumptions are not transferable. For instance, in neurons, Ca^2+^ activity is driven by depolarization-induced transmembrane Ca^2+^ fluxes. By contrast, the origins and mechanisms of astrocytes Ca^2+^ signals are diverse and still largely elusive, limiting our ability to functionally interpret Ca^2+^ events. Astrocytic [Ca^2+^]i fluctuations also primarily occur outside of the soma, within highly localized micro/nano-domains, and seldom propagate or merge at the cell body^4–7^. Concomitantly, the complexity of astrocyte morphology and its sub-diffraction limit scale also make it difficult to register areas of Ca^2+^ activity to identifiable morphological features in live fluorescence imaging. Hence, there is a growing need to enrich the astrocyte Ca^2+^ toolbox with analysis methods tailored to astrocytes. Owing to the constrains listed above, such methods need to be 1) holistic, to capture all activities in the cell, 2) unbiased, rather than restricted to user-defined regions of interest, 3) agnostic, to make no or minimal assumptions regarding the rules of Ca^2+^ propagation or integration, and 4) with instrument-limited accuracy, to favor detailed investigations consistent with the nanoscale nature of astrocytes architecture^8^.

Here, we introduce STARDUST, a pipeline that captures Ca^2+^ dynamics in confined, local micro-domains across the territory of all astrocytes in 2-photon imaging recordings. STARDUST builds upon AQuA, a popular open-source platform^9^, to yield maps of regions of activity (ROAs) from patches of active pixels, which can be combined with cell-segmentation and/or correlated to cellular morphology. Importantly, STARDUST makes no assumption regarding Ca^2+^ propagation across ROAs, in line with the seemingly static nature of astrocyte Ca^2+^ activity^4–7^. Instead, STARDUST treats ROAs as independent units and focuses on decomposing Ca^2+^ dynamics in a regionalized fashion, yielding as many as 30 ROAs per cell, or thousands across a 400 x 400 µm^2^ field of view, and extracting fluorescence time-series, signals and signal features from each of them. A particular instantiation of the usefulness of STARDUST is in pharmacology experiments, where it can distinguish “stable ROAs” (active throughout the recording), from “ON ROAs” (inactive at baseline epoch and turned on during drug application), and “OFF ROAs” (active at baseline and turned off during drug application). Together, this makes STARDUST a user-friendly complement or alternative to the small number of publicly available algorithms and tools recently developed^4,6,9–11^ to tackle astrocyte Ca^2+^ activity. This protocol includes step-by-step guidelines, tips on how to use STARDUST and outlines multiple output examples, providing a systematic walkthrough of its core functionalities, limitation, and tunability.

### Hardware

A Microsoft Windows machine with 32GB RAM is recommended. AQuA does not currently support Mac OS and Linux system.

### Software

Timing: 10 min

### Download AQuA

AQuA is a software package originally designed to decompose raw dynamic astrocyte imaging data into a set of quantifiable ‘events’^9^ based on spatio-temporal properties of voxels and assumptions that are not necessary in STARDUST. Here, AQuA is only used to extract pixels with above-threshold fluorescence independent of any user-defined region of interest or underlying cell morphology.

1. Download AQuA from https://github.com/yu-lab-vt/AquA. Follow the instruction in the section “Download and installation” to download the “MATLAB GUI” version to a local directory.
2. Extract all files from the zip folder “AQuA-master”.

Note: To learn more about AQuA, a step-by-step user guide is available in the section “Getting started” on the AQuA GitHub page.

### Install MATLAB to run AQuA

i. 3. Download MATLAB from https://www.mathworks.com/products/matlab.html. Critical: MATLAB 2017a or later is required to run AQuA.
ii. 4. Install MATLAB. During installation, install the following toolboxes required for AQuA: curve fitting toolbox, image processing toolbox, and statistics and machine learning toolbox. Alternatively, install these toolboxes in “Add-Ons” in the “HOME” tab inside MATLAB.

### Download ImageJ/Fiji for image processing

i. 5. Download ImageJ/Fiji from https://fiji.sc/.

### Install Anaconda

The STARDUST code for signal detection in Part 6 is written in Python. There are several ways to run the STARDUST code. Here we provide instructions using Jupyter Notebook through Anaconda navigator. Anaconda is a Python environment that consists of an interpreter and several installed packages. Jupyter Notebook is a web application which facilitates creating and sharing documents containing live code.

i. 6. Download Anaconda from https://www.anaconda.com/download. When clicking the download button, choose the installer version that matches your operating system.
ii. 7. Launch the installer and follow instructions to install Anaconda.

Note: The current version of Anaconda contains Python 3.11 and a fully loaded Jupyter Notebook. No additional installation for Python, Python modules or Jupyter Notebook is needed.

### Data collection

One channel image stack in tiff format is supported by AQuA. If the raw image file is in a different format, open the raw image file in ImageJ and convert it to tiff by *Save as > Tiff file* in ImageJ.

### Institutional permissions

All experiments were conducted in accordance with the guideline of the Institutional Animal Use Committee of Washington University in St. Louis School of Medicine (IACUC #20180184 and 21-0372, Animal Welfare Assurance #D16-00245).

### Key resources table

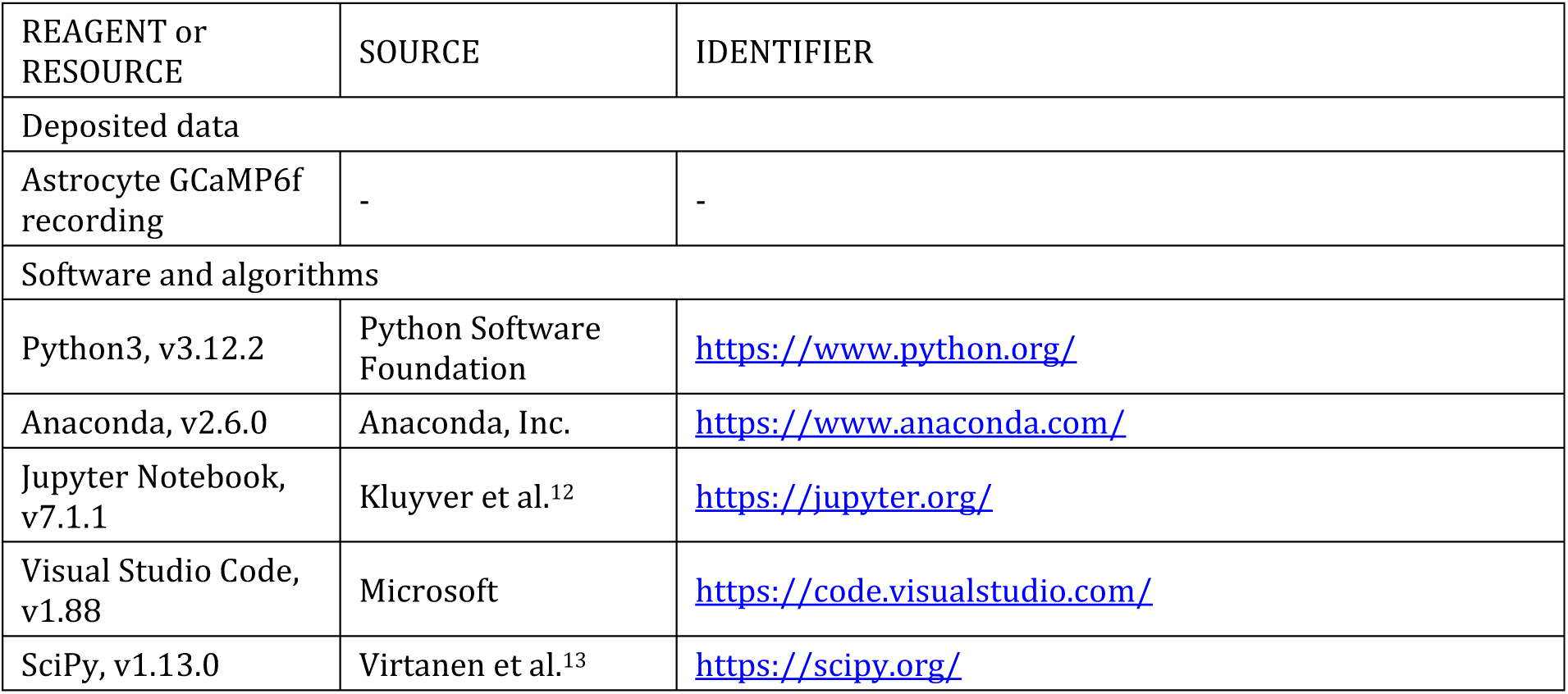
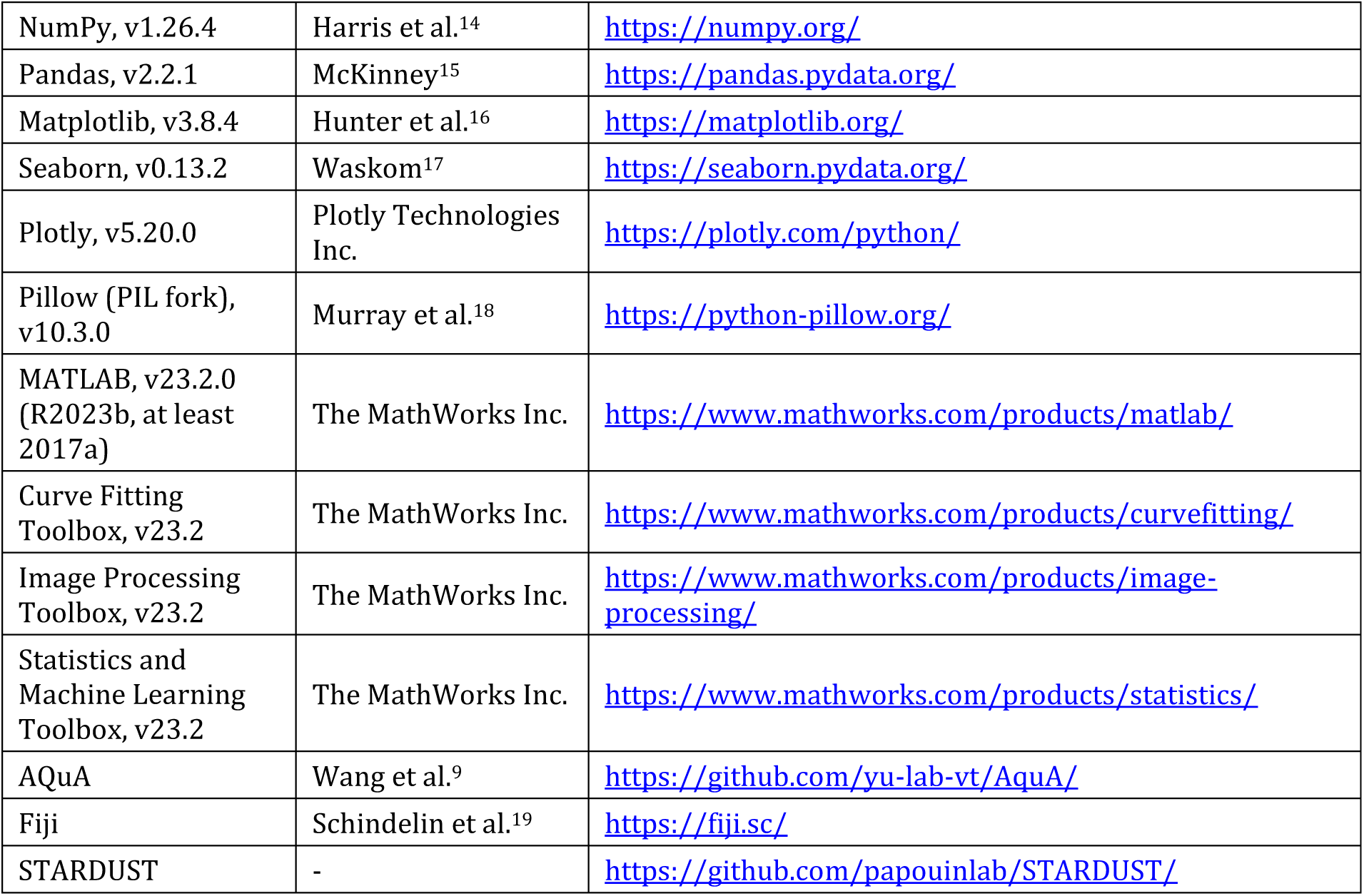

| REAGENT or RESOURCE | SOURCE | IDENTIFIER |
| --- | --- | --- |
| Deposited data |  |  |
| Astrocyte GCaMP6f recording | - | - |
| Software and algorithms |  |  |
| Python3, v3.12.2 | Python Software Foundation | https://www.python.org/ |
| Anaconda, v2.6.0 | Anaconda, Inc. | https://www.anaconda.com/ |
| Jupyter Notebook, v7.1.1 | Kluyver et al.<sup>12</sup> | https://jupyter.org/ |
| Visual Studio Code, v1.88 | Microsoft | https://code.visualstudio.com/ |
| SciPy, v1.13.0 | Virtanen et al.<sup>13</sup> | https://scipy.org/ |
| NumPy, v1.26.4 | Harris et al.<sup>14</sup> | https://numpy.org/ |
| Pandas, v2.2.1 | McKinney<sup>15</sup> | https://pandas.pydata.org/ |
| Matplotlib, v3.8.4 | Hunter et al.<sup>16</sup> | https://matplotlib.org/ |
| Seaborn, v0.13.2 | Waskom<sup>17</sup> | https://seaborn.pydata.org/ |
| Plotly, v5.20.0 | Plotly Technologies Inc. | https://plotly.com/python/ |
| Pillow (PIL fork), v10.3.0 | Murray et al.<sup>18</sup> | https://python-pillow.org/ |
| MATLAB, v23.2.0 (R2023b, at least 2017a) | The MathWorks Inc. | https://www.mathworks.com/products/matlab/ |
| Curve Fitting Toolbox, v23.2 | The MathWorks Inc. | https://www.mathworks.com/products/curvefitting/ |
| Image Processing Toolbox, v23.2 | The MathWorks Inc. | https://www.mathworks.com/products/image-processing/ |
| Statistics and Machine Learning Toolbox, v23.2 | The MathWorks Inc. | https://www.mathworks.com/products/statistics/ |
| AQuA | Wang et al.<sup>9</sup> | https://github.com/yu-lab-vt/AquA/ |
| Fiji | Schindelin et al.<sup>19</sup> | https://fiji.sc/ |
| STARDUST | - | https://github.com/papouinlab/STARDUST/ |

### Step-by-step method details

This section describes the step-by-step process for 1) image preprocessing (including motion correction and noise filtering), 2) active pixels detection using AQuA, 3) map of region-of-activity (ROA) generation, 4) time-series data extraction, 5) cell mask acquisition, and finally 6) signal detection and feature extraction with a Python pipeline.

To better illustrate each step, we use two-photon laser scanning microscopy (2-PLSM) recordings of the calcium sensor lck-GCaMP6f and static tdTomato reference marker expressed in astrocytes (AAV5-gfaABC1D::lck-GCaMP6f, AAV5-gfaABC1D::tdTomato, micro-injected at P70) in acute hippocampal slices obtained from adult mice.

Note: The goal of STARDUST is to analyze regional activity in astrocytes (**Fig.1**). If the analysis focuses on the response from the entire cell (for instance, in response to norepinephrine), the steps for detecting active pixels and creating map of ROAs should be skipped. Instead, the analysis should be as follow: 1) Part 1: Image preprocessing, 2) Part 5: Cell mask acquisition, 3) Part 4: Time-series data acquisition from cell selection, and 4) Part 6: Signal detection, steps 26-29, steps 31-35, and step 40, such that “cells” should be treated as “ROAs”. The only required input file for signal detection is the time-series data.csv file.

**Figure 1:**
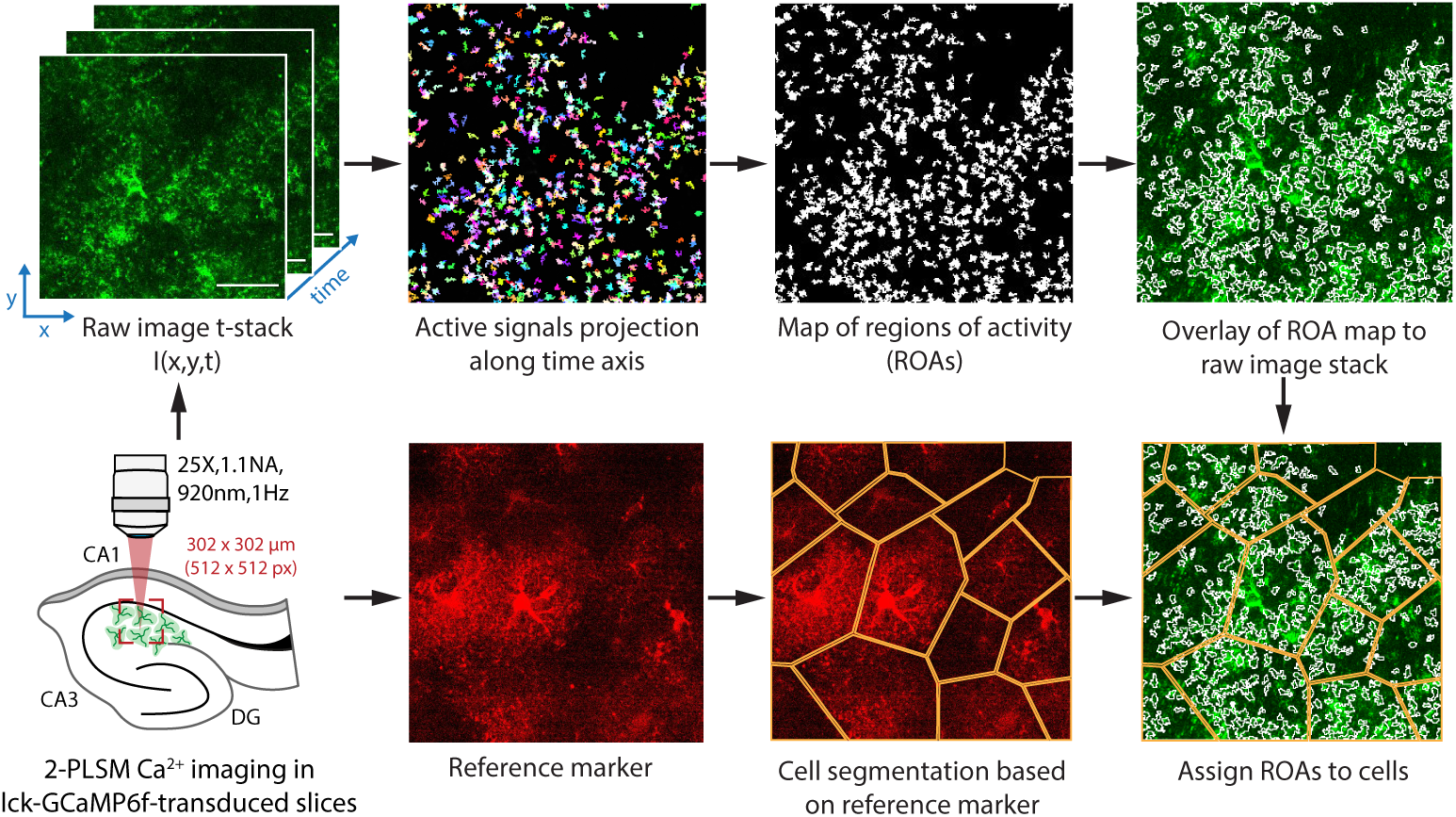
STARDUST workflow. Illustration of the STARDUST pipeline sequence. Note that cell segmentation (bottom) is optional.

### Part 1: Image Preprocessing

Timing: 5 min

This step includes recording quality check and motion correction. Because astrocyte Ca^2+^ activity occurs predominantly in small nano/micro-domains^4–7^ that are both confined and difficult to tie to any anatomically defined sub-cellular structure, astrocyte Ca^2+^ recordings and subsequent analysis do not tolerate spatial drift. We thus strongly recommend applying a motion correction against noticeable drift in the recording. If permitted, a second-channel recording of a static cell marker (e.g. tdTomato) can be used as a reference for cell registration.

1. Open the tiff image file in ImageJ.
2. Play the image stack and check the following:
3. Recording quality.
4. Open *B&C window* from *Image > Adjust > Brightness and Contrast.* Click “Auto” or adjust the slide bars to tune the contrast. A good recording should have clear and bright signals that stand out from the background noise.
5. Shift during recording and registration.
6. Identify a marker, such as a cell body or a blood vessel, as a reference to check x,y drift. If there is significant drift over time, i.e. the x,y location of signals or cell markers shift away from their original position, the recording will require a registration. Compare the last frame to the first frame to assess the extent and direction of the drift. If the recording has a significant x,y shift, TurboReg is a great ImageJ plugin for motion correction. It can be downloaded from https://bigwww.epfl.ch/thevenaz/turboreg/. Alternatively, NoRMCorre (https://github.com/flatironinstitute/NoRMCorre) is a MATLAB-based tool for non-rigid motion correction of calcium imaging data. However, due to the highly diverse nature of calcium activity in astrocytes, these tools occasionally fail to achieve an adequate registration. Therefore, the registered recording should be carefully examined before proceeding to next step. Overall, an

excessive x,y drift means that the data might need to be discarded from further analysis.

a. ii. Proceed similarly to assess vertical drift (z-axis). A noticeable shift along the z-axis means that the fluorescence data are obtained from inconsistent focal plans over time. Since this cannot be corrected, we suggest removing the recording from analysis. A significant z-shift is one where initial anatomical features (soma, branches, blood vessel) are lost from the field of view over time.
b. 3. Check the recording to assess overall fluorescence change over time, such as increased or decreased fluorescence/background activity or fluorescence bleaching: *Image > Stacks > Plot Z-axis Profile*. In the absence of pharmacological manipulations, this might be indicative of a poor recording quality. On the other hand, this is particularly useful when pharmacological treatments were performed as this might facilitate pinpointing the time of drug wash-in.

### Part 2: Active pixels detection using AQuA

Timing: 30 min – 45 min

This section uses AQuA, a publicly available software package, to identify active pixels that are above noise threshold. All supra-threshold active pixel will be color-labeled after analysis.

a. 4. Launch MATLAB. In the “Current Folder” toolbar, navigate to the folder where the file “AQuA-master” is saved. Add the “AQuA-master” folder to the working

directory by right clicking the folder name in the “current folder” window, select “Add to Path” and “Selected Folders and Subfolders”.

a. 5. Launch AQuA by entering “aqua_gui” in the Command window. A popup window of AQuA with “New project” and “Load existing” will appear. Click “New project” and a field for file selection and parameter options will show as in **Fig. 2A**.
b. 6. Input the image file to analyze by clicking the three dots on the right.
c. 7. Preset analysis parameters are determined according to the Data type selected. In the “Data type” drop-down menu, select the type of imaging condition in the corresponding data file. In our example images, data are collected from acute brain slices with lck-GCaMP6f calcium indicator, and we choose

**Figure 2:**
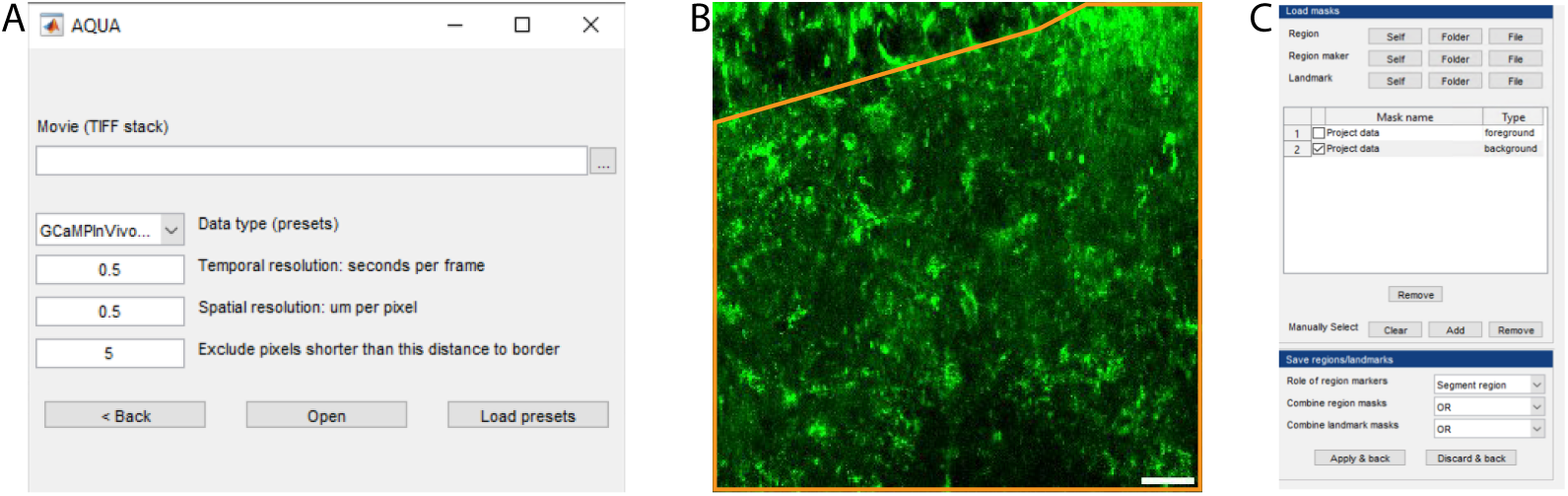
Detecting active voxels using AQuA. (A) The AQuA launch window. (B) An area of interest for subsequent analysis is drawn (orange) on the image stack reference. Scale bar: 20µm. (C) The mask builder interface.

“GCaMPExVivoLck”.

a. 8. After selecting the data type, pre-selected values will be automatically filled for the three relevant parameters at the bottom. Edit the temporal resolution and spatial resolution according to recording conditions. For “Exclude pixels shorter

than this distance to border”, change to “**0**” to ensure that the output file has the exact same canvas size as the raw file.

Critical: Setting exclude pixels at 0 is critical for accurately overlay the map of ROA back onto the raw file in Part4 step 18.

a. 9. Click “Open” to proceed to the analysis window.

Optional: Remove untargeted areas in the field of view in AQuA analysis. The top left box “Direction, regions, landmarks” includes functions such as defining cell boundary or areas or creating anatomical mask. This step can be applied to exclude undesired areas of recording or restrict the analysis to areas of interest. Specifying areas of interest reduces the processing time in AQuA by removing undesired pixels.

Here we provide an example of how to select a specific area for analysis. In this example, we are interested in analyzing the area outlined in yellow (**Fig. 2B**) and would like to exclude other areas from further analysis.

Select “Mask builder”, then in the pop-up window, select “Self” under “Region” (**Fig. 2C**). “Self” means the selected recording will be used as a reference to build the mask. After clicking “Self”, some areas are automatically selected if the pixels are above a default intensity threshold. To build a new mask that outlines the area of interest, clear the default areas by clicking “Clear” and then click “Add” to draw the area of interest by hand under the “Manually Select” section. And then click “Apply & back”. If successfully selected, the movie in the AQuA main window will have the area(s) outlined and numbered. Please refer to the AQuA user guide for other ways to determine desired areas of interest in the field of view.

a. 10. Set the parameters for detection.
b. In the detection window on the left, check the box “skip step 2,3,4” to detect active voxels without further processing and merging.

Critical: When including steps 2,3, and 4, the full-length AQuA platform assigns active voxels that coherently occur in a spatially connected regions into “events” based on their spatio-temporal properties and assumptions about event propagation that are not necessary in STARDUST. Instead, the STARDUST pipeline analyzes raw active pixels.

a. b. Set intensity threshold scaling factor, smoothing, and minimum size.

Default setting provides a good starting point for parameter selections. However, to ensure adequate detection with low false detection rate, it is crucial to optimize parameters based on recording properties, such as signal to noise ratio. For example, the intensity scaling factor can be fine-

tuned by comparing the true signals in the input recording with the color-labeled signals in the output. Increase the scaling factor if excessive noise is detected and decrease it if true signals are omitted. Minimum size can be determined by finding the smallest pixel groups that give meaningful signals, i.e. that exhibit real signals, not noise, in the raw images.

In our analysis, we use an intensity threshold scaling factor of 2.5, smoothing (sigma) 0.5, and minimum size 10 for processing. Please refer to the AQuA manual for more detailed explanation of these parameters.

a. 11. Click “Run all steps” to run the pipeline. A proof-reading window and export window will appear once the software finishes processing.

Note: The processing time depends on the size, quality of the recording, and computer hardware. For a 600s recording with a field of view of ∼400µm^2^, the processing time is ∼30 minutes.

a. 12. Set the filter condition for the output file. This step is crucial for Part 3.
b. Select the parameter “Area”, which is the size of the active zones in µm^2^, as the filter criteria by checking the adjacent box in the “Proof reading” window (**Fig. 3A**). The range of area needs to be optimized to capture as many regions of activity (ROAs) and signals as possible in later steps. We strongly recommend testing out different ranges of area min and max cutoffs because the adequate range will depend greatly on the imaging conditions and image stack quality (see **Fig. 5** below). Too stringent a filter criterion will lead to the loss of active regions, while too loose a criterion will combine/merge adjacent regions into a single ROA leading to a loss of spatial resolution and decrease in signal to noise ratio as illustrated in **Fig. 3** (<u>Troubleshooting 1</u>).

**Figure 3:**
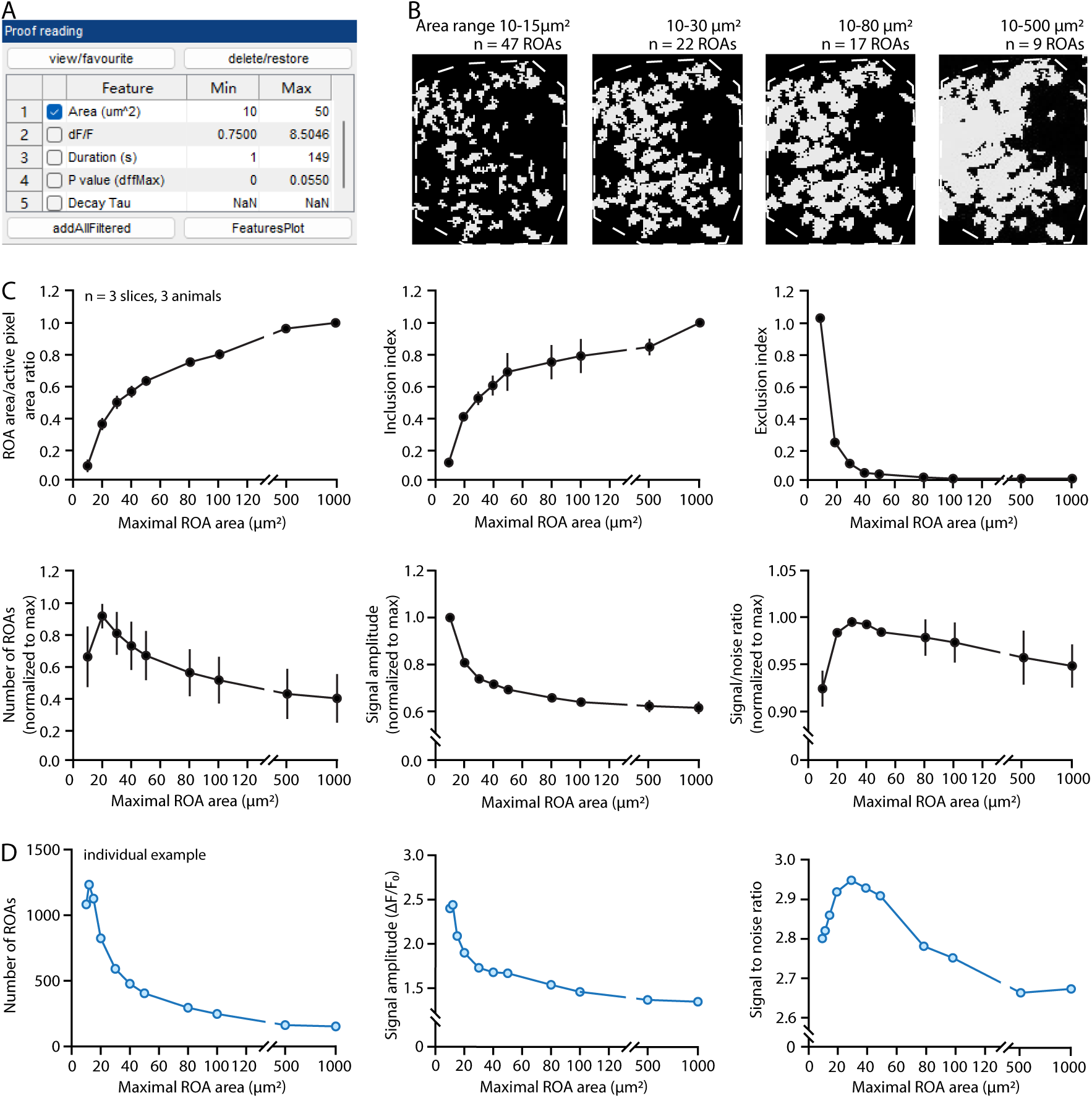
Determining the adequate range of ROA sizes. (A) “Proof reading” window in AQuA for selecting appropriate filter criteria of the output movie. (B) Effect of the min and max cutoff values for the “Area” parameter on the map of active zones (greyed areas) over a representative astrocyte (cell boundary is denoted with dotted outline). The number of ROAs eventually obtained with each max area cutoff is indicated. Note how, as active zones get larger, they tend to merge, resulting in larger but fewer ROAs (also see below). Recording was acquired on Nikon A1R microscope. (C) Effect of the max Area cutoff on diverse parameters. The inclusion index reflects how the map of ROAs, for a given max area cutoff value, successfully captures the large active pixel patches that might be excluded by this setting. It is measured as the percentage of area of the excluded large pixel patches that overlaps with the final map of ROAs. The exclusion index measures the number of active pixel patches that have no pixels included in the final map of ROAs. Data are shown as mean ± sem. Recording was acquired on Nikon A1R microscope. (D) Example of the effect of max area cutoff on the final number of ROAs, average signal amplitude and signal to noise ratio for an individual experiment. Note how peaks, optima, or minima may occur for different max area values across experiments (compared to C). Recording was acquired on Nikon A1R microscope.

Note: We recommend keeping the max area cutoff value within a narrow range (in our case 20µm^2^ to 40µm^2^) across the same dataset but adjusting it as needed for each individual recording. The min area cutoff value is usually set the same as the detection setting.

a. b. Export the movie with color-labeled active signals overlay. In “Movie brightness/contrast” window on the right, set the brightness of the raw

recording to zero, and in the “Feature overlay” window, set the brightness of the overlay image to max.

a. 13. Save the output file. In the export window on the left, uncheck the “Events” and “Feature table” box, leaving only the “Movie with overlay” box checked. Click “Export/Save” to save the file in the targeted folder.

Optional: Save the processed AQuA file. Select “Events” in the export window and save to directory. Saving the AQuA file allows for adjusting filtering criteria and output new overlay movies later if needed. To access a saved AQuA MATLAB file, select “Load existing” after starting an aqua_gui window in step 2.

### Part 3: Acquisition of the map of regions of activity (ROAs)

Timing: 5 min

This section described how to generate the map of regions of activity (ROAs) from the AQuA output movie using ImageJ. We define an ROA as the aggregate spatial footprint formed by the t-projection of a confined ensemble of active pixel patches. As a result, 1) an ROA is a spatial domain within which at least one active pixel patch occurs during a recording, 2) active pixel patches within an ROA do not need to occupy the entire spatial footprint of the ROA, 3) the footprints of two active pixel patches from a same ROA do not necessarily overlap, as it is the aggregate footprint of all active pixel patches that defines the boundaries of the ROA, 4) any two given ROAs are spatially unconnected, and 5) the constellation of ROAs of a given cell represents a map of spatially independent zones of activity within that cell. The activity of two contiguous ROAs is assumed to be independent; however, occasionally, large active pixel patches will be excluded from the map of ROA acquisition process, due to the maximum area cut-off criteria, and may result in concomitant signals in several contiguous ROAs. In total, the advantages of this analysis method are that it is unbiased, semi-automated, comprehensive, and solely activity-based in that it is decoupled from (but compatible with) morphological constrains or cell-based criteria.

a. 14. Acquire the map of ROA by performing a t(z) projection on the output file from AQuA. Open the file in ImageJ. Click *Image > Stacks > Z Project > Max intensity* to generate a t-projected image.
b. 15. Assign ROI numbers to each ROA for analysis in ImageJ.
c. Convert the t-projected ROA map to binary image by *Process > Binary > Make Binary*.

Note: ImageJ does not have an “ROA” manager, instead we use the ROI manager for any selected regions of interest, whether ROAs or cells (see Part 5). Beware of possible confusions in the subsequent text.

a. b. Then select *Analyze > Analyze particles*, set the size to 0-infinity. Also check the boxes of “Include holes”, “Add to manager” and “Composite ROIs”. Click “OK”.

Note: In the ROI manager, each ROA is assigned a number and highlighted with colored boundary in the image. The total number of the ROAs can be found as the last number in the ROI manager.

Note: Large, connected ROAs may arise if the filtering criteria in step 12 are not optimized (Troubleshooting 1).

a. 16. Save all the ROAs (ROI selections). In the ROI manager, select all ROAs and right click to save as a ROA_RoiSet.zip file. This file can be reloaded into ImageJ if the ROA map needs to be adjusted later.
b. 17. Generate a binary ROA mask.
c. Select all the ROAs in the ROI manager (Ctrl+A), right click and select “OR (Combine)”, then click “Add [t]” in the ROI manager. This generates a new ROI that combines all the ROAs.
d. Select this ROI and then go to *Edit > Selection > Create Mask*. A binary mask with all of ROAs shown as white area is generated. Save this mask under the same file folder as all other files by clicking *File > Save as > Tiff* file.

### Part 4: Time-series data acquisition

Timing: 3 min

This section extracts fluorescence intensity time-series from each ROA in the ROA map.

a. 18. In ImageJ, open the original raw recording tiff file. Open the ROA_RoiSet.zip file generated from step 16. Overlay the ROA selections onto the original file by clicking “Show All” in the ROI manager.
b. 19. Set measurement item in *Analyze > Set measurements*. Check the box “mean gray value”. This measures the intensity of each selected ROA.

Note: The area of each ROA can be obtained with the same method by checking the box “Area” instead. This is of interest when comparing ROA features (e.g., numbers, size) across different conditions (see **Fig. 5** for example).

a. 20. Select all the ROAs the ROI manager, right click and select “multi measure”. In the next popup window, select “measure all slices” and “one row per slice”. A result window will then appear with the values of intensity for each ROA over time.
b. 21. Copy and paste the data from the result window into an excel file. Save as csv file format under the same folder where the ROA mask is.

**Figure 4:**
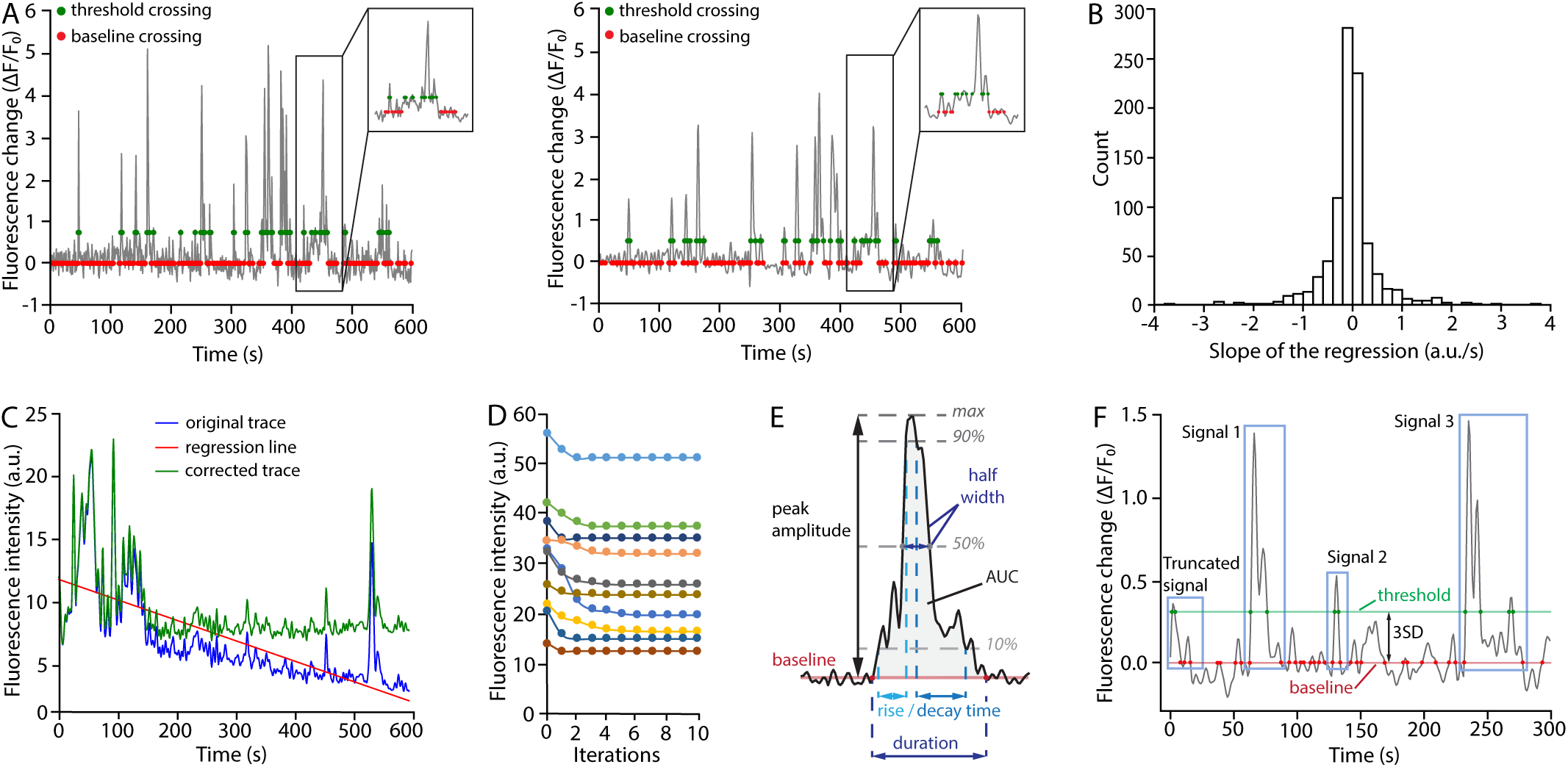
Signal detection and analysis. (A) Left: raw ΔF/F0 trace; Right: ΔF/F0 trace filtered with a Butterworth filter. Note how after filtering the amplitude of signals is visibly diminished. Due to noise reduction, however, the 3SD threshold is also significantly lowered, resulting to minimal or no changes in signal detection. However, signal features might be affected. Red dots indicate where the traces crosse F0. Green dots indicate where the traces crosse the signal detection threshold. (B) Distribution of the linear regression slope from all ROA traces from a single recording. In a stable recording like this one (no or minima drift), the distribution is centered near 0 (see Fig. 7A for opposite example). (C) Example of drift correction. The downward drift in the original trace (blue) is corrected by subtracting a linear trend (red line). The corrected trace (green) is used for subsequent analysis. (D) Baseline fluorescence intensity (F0) of 10 individual time series over 10 cycles of iterations showing the rapid convergence of the baseline fluorescence determination process. In our hands, the baseline fluorescence value systematically converges within 5 iterations (usually 3). (E) Schematic illustration of the main signal features and how they are measured. AUC: area under the curve; max: peak signal amplitude from baseline. (F) Illustration of a fluorescence time series extracted from a single ROA, showing baseline crossing points (red dots) and signal detection threshold crossing points (green dots) for 3 signals. Note the presence of a ’truncated’ signal due to the absence of an initiation point, which will be excluded from further analysis. 3SD: 3 standard deviations.

**Figure 5:**
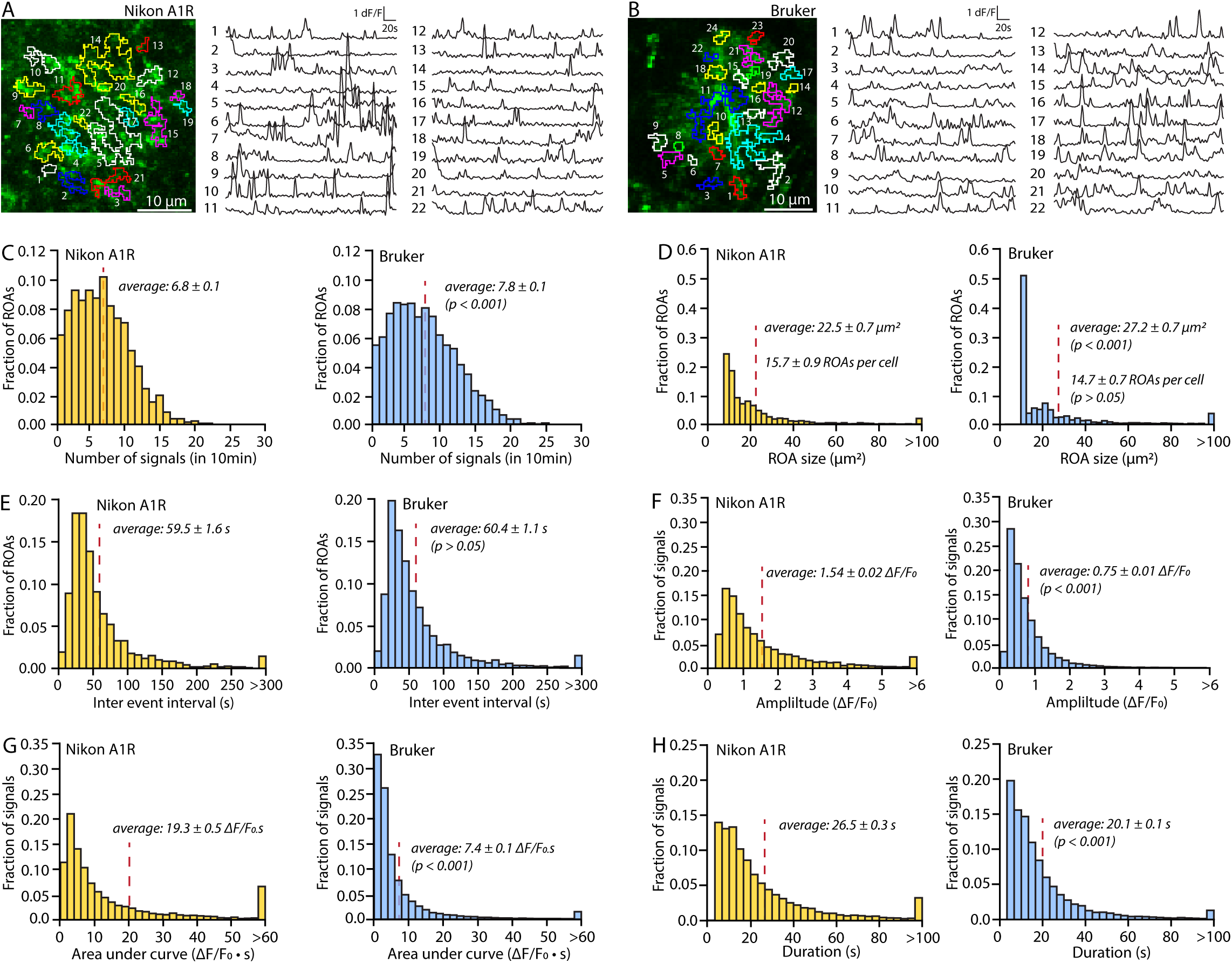
STARDUST benchmarking. (A and B) Representative maps of ROAs overlayed on a maximum projection of an individual astrocyte, and fluorescence time series extracted from each ROA, for experiments conducted with a Nikon A1R (A) and a Bruker Ultima 2pPlus (B). Imaging conditions were the same otherwise (25X, 1.10NA, 37.5mW, recording rate 1Hz). (C - E) Histograms illustrating the number of signals (C), ROA size (D), and inter-event interval distributions (E) across ROAs in 10min recordings acquired with a Nikon A1R and Bruker 2-PLSM. Permutation tests were used, n=3 slices from 2 mice. (F - H) Histograms illustrating the amplitude (F), area under the curve (D), and duration distributions (F) of signals detected in 10min recordings acquired with a Nikon A1R and Bruker 2-PLSM. Permutation tests were used, n=3 slices from 2 mice.

**Figure 6:**
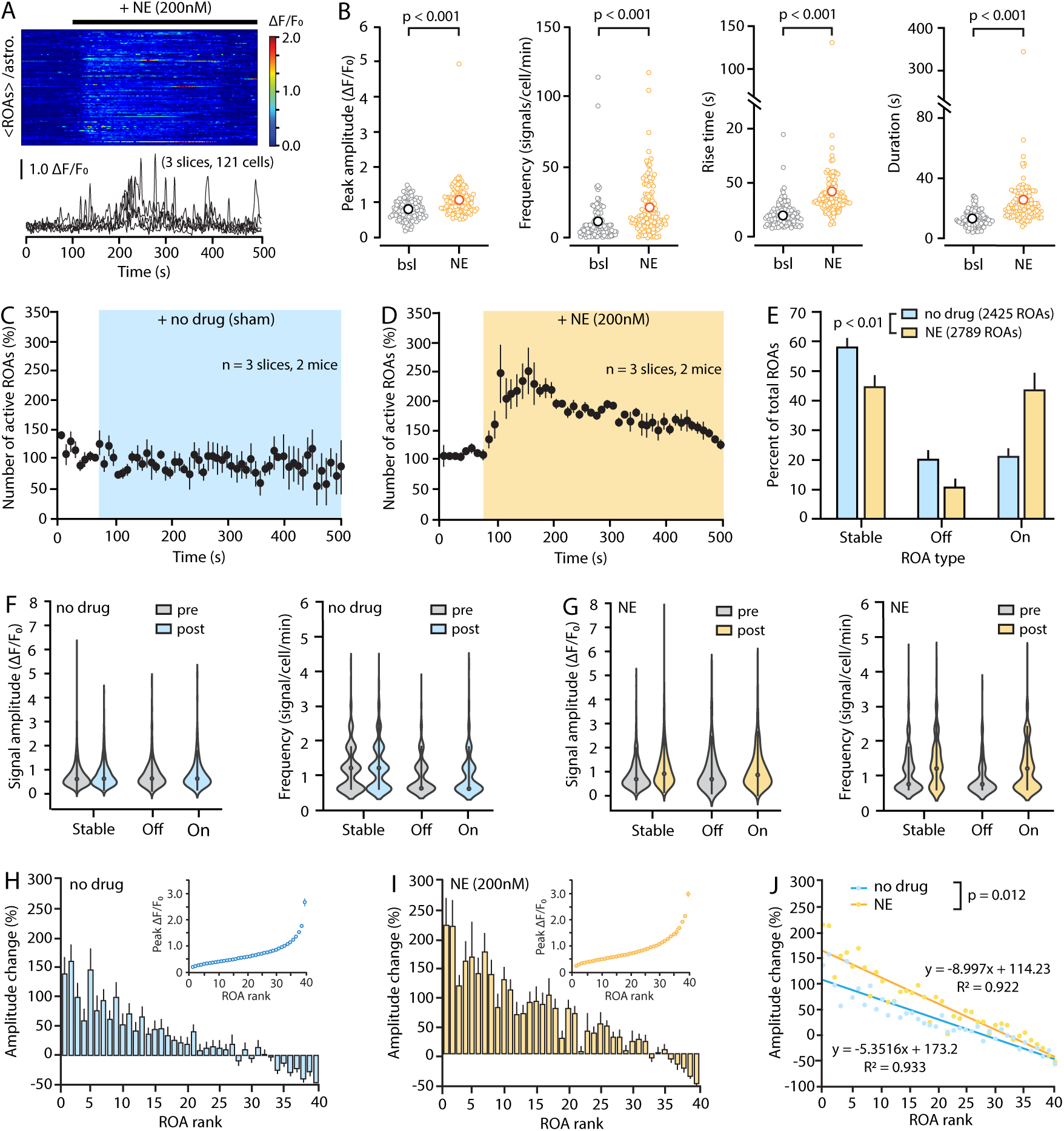
ROA-based analyses of calcium responses to norepinephrine using STARDUST. (A) Kymograph (individual rows show averaged ROA fluorescence per astrocyte) and 5 representative ΔF/F0 traces (± sem) of spontaneous astrocyte Ca2+ activity in response to the bath application of a ‘subthreshold’ concentration of NE (200nM, i.e. causing no cell-wide response). (B) Plots showing the effect of 200nM NE on the amplitude, frequency and kinetics of Ca2+ signals compared to the baseline epoch (bsl). Data shown as averages of all ROAs per cell. Permutation tests were used. (C and D) Time courses of the effect of a no-drug sham application (C) and 200nM NE bath application (D) on the number of active ROAs (10s bins) normalized to baseline. (E) Proportions of “Stable”, “On”, and “Off” ROAs for the experiments shown in (C) and (D) (no-drug vs. NE: p = 0.0019, Chi-square test). (F and G) Signal amplitude and frequency per ROA type in no-drug (F) and NE (G) conditions. Effect of ‘no-drug’ on amplitudes across ROA types: p>0.05, linear mixed effect model. Effect of ‘no-drug’ on frequency: Stable-pre vs. Off p<0.001, vs. On p<0.001, Stable-post vs. Off p<0.001, vs. On p<0.001. Effect of NE on amplitude: Stable-pre vs. post p<0.001, vs. Off p<0.001, vs. On p<0.001. Effect of NE on frequency across ROA types: p<0.001, linear mixed effect model. (H and I) ROA-ranked based analysis of the effect no-drug (H) and NE (I). ROAs are ranked based on the peak signal amplitude detected in the corresponding time-series during the baseline epoch (inset, 40 bins). The percentage change in amplitude after drug application is then plotted as a function of ROA rank. (J) Linear regression analysis of the experiments shown in H) and I). NE exerts a differential effect across ROA ranks that is greater than that observed by ‘chance’ (NE vs. No-drug, p = 0.012, combined regression model). Data are obtained from 100s of pre- drug condition (0-100s) and 100s of post-drug condition (200-300s) from stable ROAs.

### Part 5: Cell mask acquisition

Timing: 30min

This step identifies the cell boundaries in the field of view. This step is optional for ROA-based analysis. However, it allows assigning ROAs to individual cells, which permits cell-based averaging of ROA data. It is also necessary for cell-based analysis, i.e., if the fluorescence time series need to be extracted from whole cells.

a. 22. Obtain a reference image for cell boundary in the recording field of view. Because of the complex structure of astrocytes, determining the cell boundary for each cell is intrinsically challenging. We offer two suggestions to obtain a reference image.
b. If multichannel recordings or images from a second, static marker were acquired: the reference image can be obtained from them. In our analysis, tdTomato, used as a call marker, and lck-GCaMP6f are co-expressed in astrocytes and imaged in the red and green channel respectively.
c. If no static markers were used: a reference image can be obtained by generating a z-projection from the original t-stack recording by *Image > Stacks > z projection*. Because baseline fluorescence in the lck-GCaMP6f is usually low, and because cyto-GCaMP6f primarily reveals astrocytes soma and branches, the arborization of individual astrocytes may not appear in full in either case. However, this might suffice to outline the general shape of astrocytes.
d. If cell segmentation is of paramount importance, we strongly recommend using sparse expression of GCaMP6f (usually by reducing viral titer or tamoxifen dosage) and other cell segmentation tools such as Imaris.
e. 23. In ImageJ, open the reference image and the ROA_RoiSet.zip file, and overlay the ROA selections onto the reference image by clicking “Show All”. Delineate cell boundaries with the “Polygon selections” tool making sure to include each ROA in its respective cell. Add each cell selection to the ROI manager by “Ctrl +t” or “Add” in the ROI manager.

Note: All ROAs should be captured within a cell boundary, since astrocytes tile the entire neuropil. Additionally, because astrocyte domains have virtually no or minimal spatial overlap, cell boundaries should not touch each other <u>(Troubleshooting 2).</u> Note that, by virtue of their 3D tiling, it can be challenging to determine whether a set of processes/ROAs belongs to an adjacent astrocyte within the focal plan or to an astrocyte outside of the focal plan. Here again, if this is an issue, see note 22.b. and 22.c. above.

1. 24. Select and delete all the ROAs from ROI manager, leaving only the cell selections in the ROI manager. Select all and right click to save as a CELL_RoiSet.zip file.
2. 25. Generate the cell mask.
3. Select all the cell selections in the ROI manager, right click and select “OR (Combine)”, then “Add” to the ROI manager. This generates an ROI that includes all the cell selections.

Critical: Overlapping cells will be recognized as one continuous cell in later steps. Visually verify that there is no overlap between cells. Adjust cell boundaries if necessary.

1. b. Select this ROI and then go to *Edit > Selection> Create Mask*. Save it as Cell_mask.tiff file under the same file folder. This folder should now contain the cell mask tiff file, the ROA mask tiff file, and the time series csv file.

Note: To this point, the ROA mask and cell mask as tiff files and the fluorescence time-series data stored in a csv file have all been generated. These are the three input files needed for downstream signal detection and analysis with the STARDUST Python pipeline.

### Part 6: Signal detection and feature extraction

This section introduces an interactive and user-friendly Jupyter Notebook Python script for signal detection and analysis, which includes signal preprocessing, baseline fluorescence (F0) determination and signal detection, signal feature extraction, ROA-based averaging and data output. When running STARDUST, the script will take users through the pipeline semi-automatically, with interactive prompts, stopping at steps that require user inputs or provide output graphs for visual inspection. To streamline the process, the Jupyter Notebook is equipped with custom-defined functions stored in util.py. All scripts are available on https://github.com/papouinlab/STARDUST.

This section provides explanation for the inputs required by each code block, the underlying computation, and the outputs they give. For more detailed explanation of arguments and output, users can use the help documentation for each function by executing help(***function_name***) in the interactive environment. Notes on how to optimize key parameters are also detailed in this section.

Note: Code segments shown in this section are provided in the “STARDUST

interactive.ipynb” file in the GitHub repository. Unless explicitly noted, the users do not need to edit the content of the code.

Note: Code blocks are referred to as “cells” in the Jupyter Notebook ecosystem. To avoid confusion, we will be referring to these as code blocks in this protocol.

Note: In the STARDUST pipeline, each non-optional code block needs to be run before executing the next block. To run a code block, click anywhere inside the code block to select it, click the “Run” button on the top of the Jupyter Notebook page or use Shift + Enter to execute the code. There will be a set of empty brackets [ ] on the left side of each code block. When the code is running, the brackets will show an asterisk sign inside [*]. When the code completes running, a number will show up in the bracket, marking the order of executed code blocks.

Note: Some code blocks, upon running, will prompt users for input in a text box under the code block. Input the prompted information and press Enter to proceed to the next prompt.

1. 26. Download the STARDUST repository from GitHub. On the top right of the repository webpage, click the green “Code” button, click “Download ZIP” to a local directory. Extract all files.

Note: Because online Jupyter Notebook navigating to the C Drive in Windows system by default, we recommend downloading the Zip file into the C Drive.

1. 27. Launch Jupyter Notebook from the Anaconda navigator. This will open a browser tab showing the Notebook Dashboard. The Dashboard is similar to a file explorer. Navigate to open the STARDUST GitHub folder “STARDUST-main”, and open the STARDUST Jupyter Notebook file named “STARDUST interactive.ipynb”.

Note: If performing whole cell-based analysis, open “Cell-based STARDUST.ipynb” instead. This file only contains code blocks necessary for the whole cell-based analysis. To continue, proceed to steps 28-29, steps 31-35, and step 40.

1. 28. Execute the following code block by clicking “Run”, which will import modules and custom-defined functions.
2. 29. Run the next code block to read in input files and information of the experiment. Upon running, the code will prompt users to input the path to the folder that contains all input files, the file names, and information about the experiment including timing of pharmacology application, frame rate, and spatial resolution of the recording. Enter the information accordingly in the prompted text boxes.

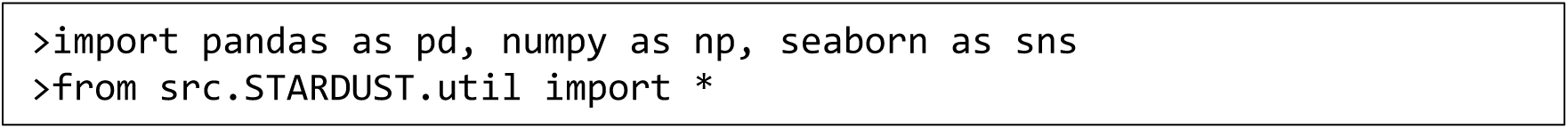

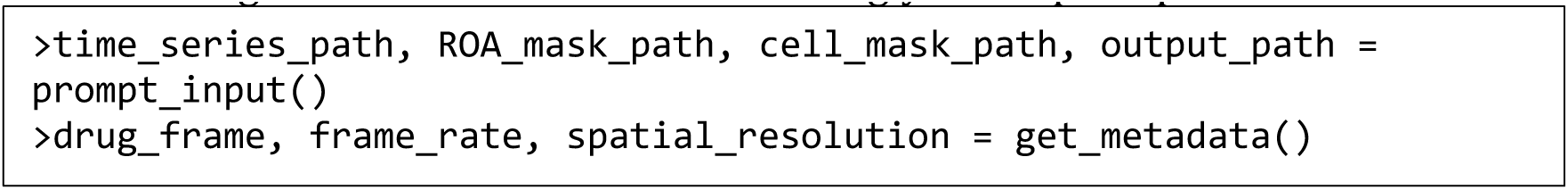

Note: To find the path to a folder in a Windows system machine, navigate to the folder, click on the address bar in the file navigator and copy the path. For example, the path to a folder named “example folder” under the Documents folder should look like “C:\Users\YourUserName\Documents\example folder”.

Critical: The input files, including the time series csv file (generated from step 21), the ROA mask tiff file (step 17), and the cell mask tiff file (step 24) must all be under the same folder.

1. 30. Run the code block below to read in the ROA mask and cell mask. This also labels ROAs and cells to allow ROA assignment to a specific cell.

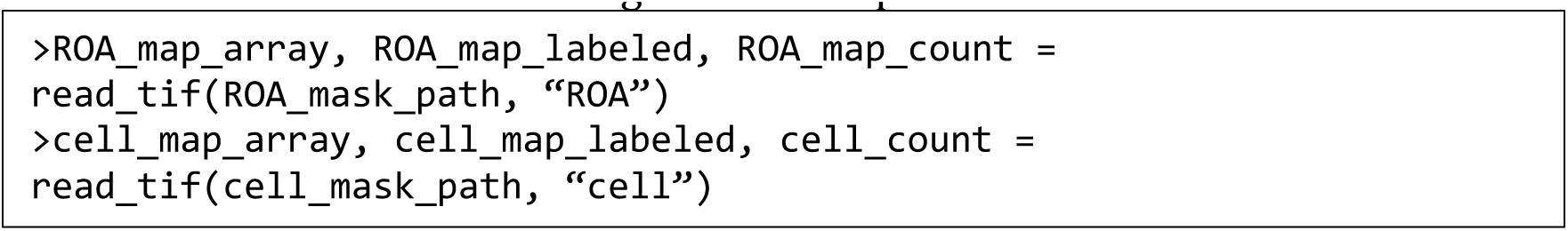

Critical: This code block outputs the total number of ROAs or cells being analyzed. The number should match the number of ROAs or cells generated in earlier steps in ImageJ (<u>Troubleshooting 2</u>).

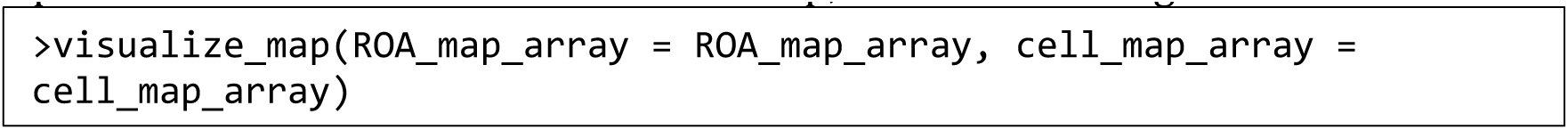

Optional: To visualize the ROA and cell map, run the following code block.

### Signal preprocessing

Timing: 2 min

This step provides signal smoothening and detrending. However, due to the enigmatic and dynamic nature of astrocyte calcium activity, signal preprocessing may alter or eliminate true features of calcium activity and therefore should be used with moderation, great caution and adequate rationale.

1. 31. Run the code block to extract raw traces from the input file and generate filtered traces.

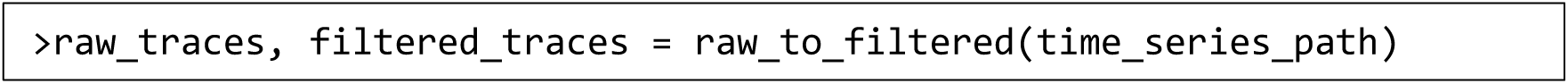

Note: To smoothen fluorescence signals, a 4^th^ order low-pass Butterworth filter with a cutoff frequency of 0.4 Hz is applied to the signal traces, which filters the signal in a forward and backward direction to remove phase distortion. The low-pass filter is set to remove the high-frequency noise, while the cutoff frequency is set to ensure a minimal modification of the traces (**Fig.4A**). The order and cutoff frequency can be adjusted by specifying optional arguments *order* and *cutoff* in the ***raw_to_filtered*** function.

Note: Users can choose to use the filtered traces or the raw traces in the following steps. By default, the filtered traces are used for further processing. To proceed with the raw traces instead, change “filtered_traces” to “raw_traces” in the code block in step 32.

1. 32. Run the ***check_traces*** function to extract and output the total ROA number and number of frames. The result should match the number of ROAs from the ROA mask and the recording length in frames.

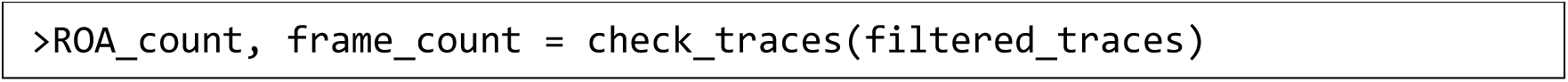

Optional: Run the ***correct_shift*** function to visualize the slope distribution of all traces obtained using linear regression and probe potential fluorescence shift **(Fig.4B**). If the slope distribution is not centered around zero, this might indicate a drift or photobleaching during the recording. This function then corrects linear trends by a default factor of 0.5 from the original traces (**Fig. 4C**).

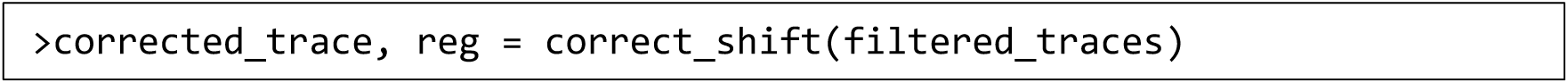

Note: The degree of correction can be adjusted using the optional argument *correction_factor* which represents the percentage of linear trend to be subtracted in the function. <u>Troubleshooting 3</u>). This function then corrects linear trends by a default factor of 0.5 from the original traces (**Fig. 4C**).

Note: Linear regression is best used to remove a trend shared by all time-series in a recording (which is an indication that an overall drift occurred during the recording). By contrast, we advise that an upward or downward drift in only a sub-selection of ROAs might be indicative of a local or cell-specific change in baseline calcium levels, which is potentially biologically relevant and therefore should not be corrected. Additionally, beware that large signals tend to bias the linear regression calculation. Lastly, there exists other sophisticated methods for signal correction. Please select the optimal methods in specific conditions.

### Baseline fluorescence determination and signal detection

Timing: 2 min

Baseline fluorescence (F0) determination is a prerequisite for accurate signal detection and features extraction. F0 is determined by iterating a three-step process: 1) interim F0 determination, 2) above-threshold segments identification, 3) segments removal. During the first iteration, F0 is calculated as the average of the entire trace. Next, trace segments above the signal threshold (i.e. potential signals) are identified and removed. The interim F0 value at subsequent iterations is thus determined as the mean of the ‘signal-free’ trace. We find that F0 rapidly converges onto a value corresponding to the mean fluorescence of an entire trace devoid of signals exceeding the selected detection threshold (**Fig.4D**).

1. 33. Run the following code block to determine F0.

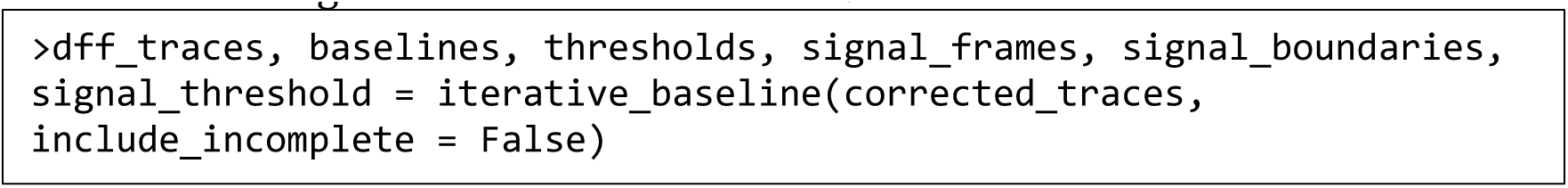

Upon running, the function will ask to input a desired number of iterations, and signal threshold. We find that convergence occurs within 5 iterations with no further improvement (and no deviation) with further iterations (**Fig.4D**). As for the signal threshold, we, and others, determine signal segments in traces based on the standard deviation (SD) in F0. We recommend setting signal threshold as 2-to- 3SD above F0. Once the F0 is determined, the change in fluorescence intensity relative to F0 is calculated for each data point (ΔF/F0 trace) as: (F(t) – F0)/ F0.

A signal is defined by “F0 crossing points”, when the trace intersects the average F0, and “threshold crossing points”, when the trace intersects the signal threshold (**Fig. 4F**). The bounds of a signal are thus the nearest F0-crossing points flanking one or more threshold-crossing points (**Fig. 4E**).

Note: Per this design, signals that lack an initiation or termination bound (i.e. one of the two F0 crossing points) will not be captured for analysis (**Fig.4F**, “Truncated signal”). This might result in some ROAs having no signals, even though they were derived from an active pixel patch. In occasions when truncated signals need to be included (for example, large signals with slow decay times that do not return to baseline during the recorded epoch) specifying the optional argument *include_incomplete* as True allows for extractions of available features (<u>Troubleshooting 4</u>).

1. 34. Run the code block to visualize all ΔF/F0 traces of the current time-series file as a heatmap (also known as kymograph). Each line will show color-coded ΔF/F0 values of a single ROA overtime.

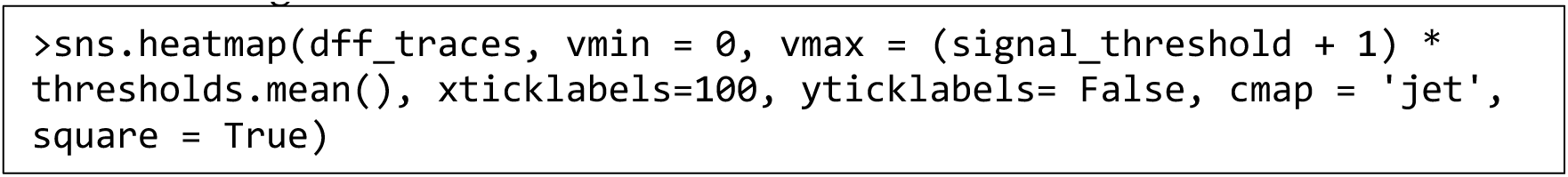

Note: To facilitate visualization, we manually specify the optional arguments *vmin* and *vmax* that set the range of the color map. *vmin* specifies the lower bound of the color map. In our case we have set *vmin* at 0, meaning that ΔF/F0 values lower than 0 will show as dark blue. Similarly, *vmax* specifies the upper bound of the color map. We choose to set *vmax* at (signal_threshold + 1) * average thresholds, so that signal segments with a ΔF/F0 value larger than *vmax* will be colored red. *vmin* and *vmax* can be adjusted as needed.

### Signal feature extraction

Timing: 30 s

1. 35. Run the code block to extract signal features with the ***analyze_signal*** function. This code generates a dataframe stored in signal_stats, with each row representing one active signal. The columns detail signal features listed in table 1 (**fig. 4E**).

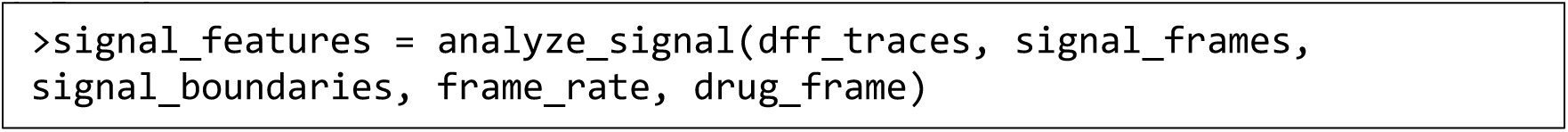

**Table 1.**
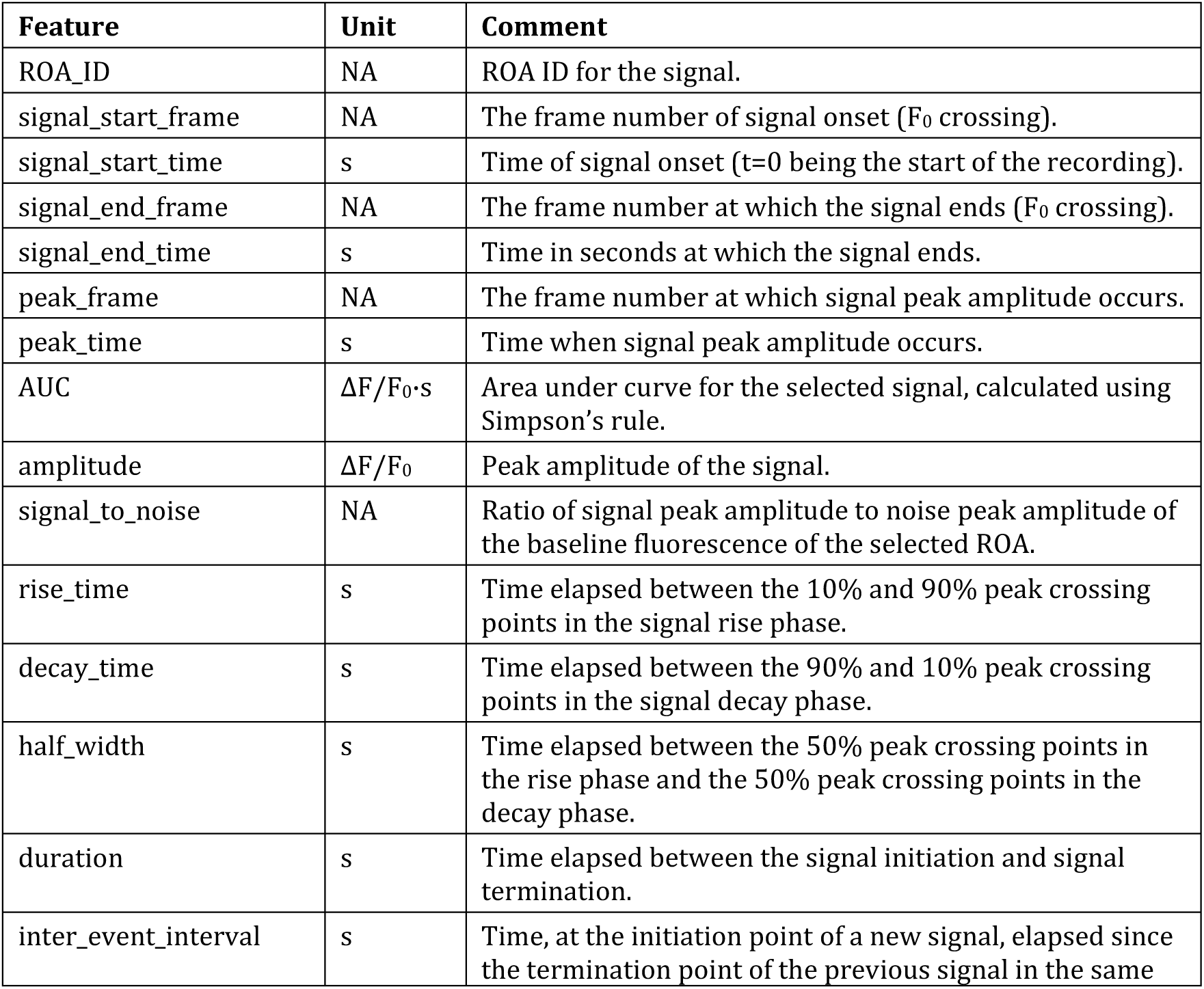
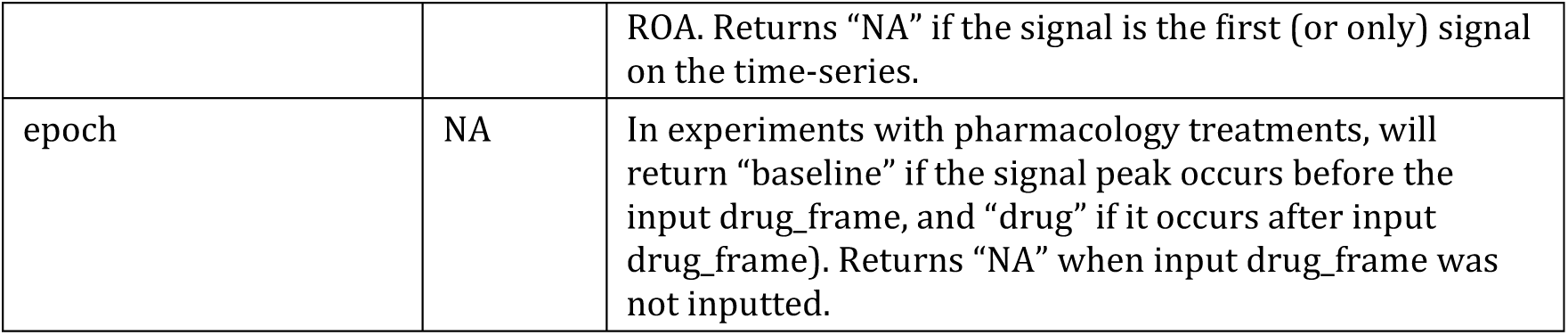
Signal features output dictionary. Note: These definitions depart slightly (but importantly) from those typically employed for neuronal recordings. For instance, in neuronal recordings, the decay time is often the time constant of the exponential fit applied to the “90%-10% of peak amplitude” portion of the trace, and the inter-event interval is determined from signal onset to signal onset. However, few astrocyte Ca^2+^ signals appear to display EPSC- or EPSP-like kinetics, and in most Ca^2+^ recordings of astrocyte activity the sampling rate (∼1Hz) is often too low for an adequate exponential regression, and the duration of astrocyte Ca^2+^ transients are highly variable, prompting this minor differences.

### ROA-based analysis

Timing: 2 min

This step uses the two binary masks-ROA_mask.tiff and Cell_mask.tiff to assign each ROA to the cell with which it overlaps. This step is applied to group ROAs of a given cell together and carry out subsequent cell-based averaging of ROA data.

1. 36. Run the ***align_ROA_cell*** function to assign the corresponding cell IDs to each ROA using the ROA and cell masks provided. The function also calculates ROA size.

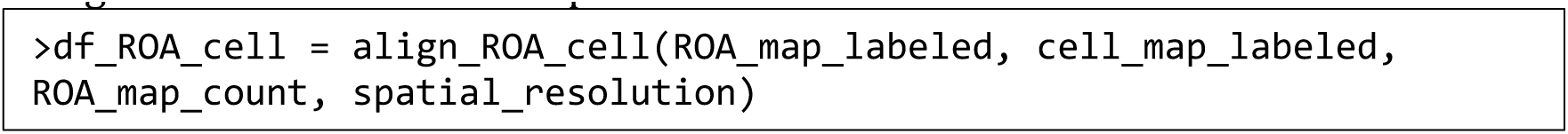

Note: ROAs that were not circled into any cells are assigned to “cell 0”.

1. 37. Run the code block to add ROA and cell information to the extracted signal feature data frame.
2. 38. Run the next code block to perform ROA-based analysis. The ROA-based analysis summarizes averaged signal features per ROA (table 1) and ROA specific features (table 2). In addition, this step also categorizes each ROA according to their activity in response to pharmacological treatments (when applicable).
3. 39. Run the next code block to generate cell-based averaging of ROA data.

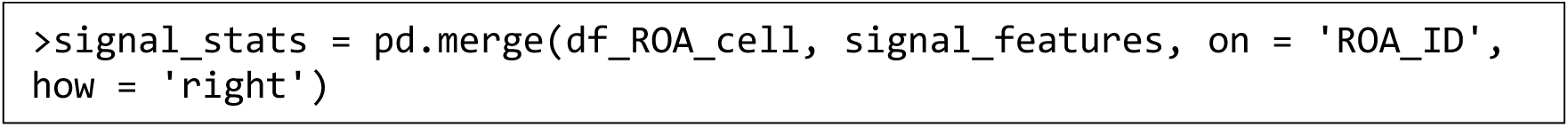

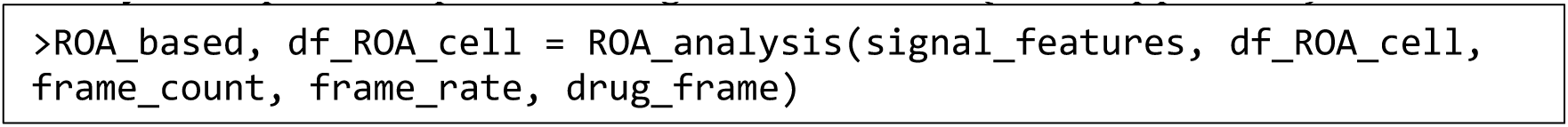

**Table 2.** ROA-based analysis output dictionary.

| Output | Unit | Comment |
| --- | --- | --- |
| ROA_ID | NA | ROA ID. |
| cell_ID | NA | Cell ID of selected ROA. |
| size_um2 | $\mu\text{m}^2$ | Size of the selected ROA. |
| ROA_type | NA | ROA categorization based on its activity with regards to the drug_frame input.<br>If signals were detected in the baseline and drug perfusion epochs of the time-series of the selected ROA, the ROA is categorized as “stable”. If the time-series contain signals during ‘baseline’ epoch but non during the ‘drug’ epoch, it is listed as an “off” ROA. If the time-series contains signals in the ‘drug’ epoch but none in the ‘baseline’ epoch, it is listed as “on” ROA. If no signals are detected, the ROA is listed as “inactive” (See Troubleshooting 5 if a recording contains a high proportion of “inactive” ROAs). Returns “NA” if no value was inputted for drug_frame. |
| epoch | NA | Epoch from which average measurements were obtained. |
| signal_count | NA | Number of signals in the indicated epoch. |
| frequency_permin | $\text{min}^{-1}$ | Number of signals per minute in the selected ROA during the indicated epoch. |

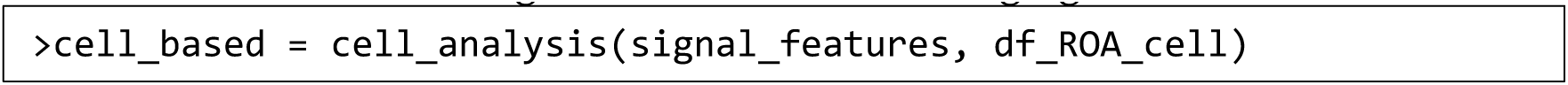

### Data output

Timing: 30 s

1. 40. Run the next code block to output the analysis results. The results are compiled as pandas dataframes and can be exported as an Excel file with multiple datasheets or separate csv files by specifying the *save_as* argument as “excel” or “csv”.

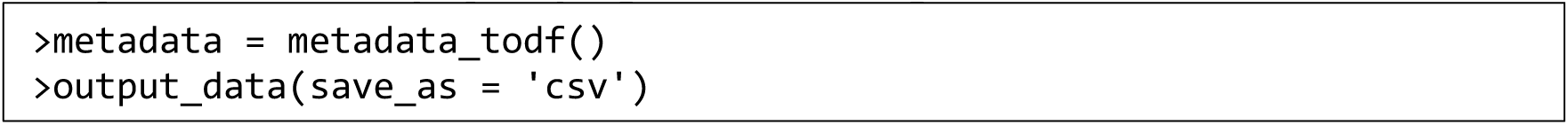

Depending on the analysis results, the outputs include, but are not limited to, metadata containing important parameters such as the time of drug application, the list of ΔF/F0 traces for all ROAs, the list of ΔF/F0 traces averaged from ROAs per cell, the master sheet with all features analyzed for each signal, the master sheet for ROA-based analysis and the cell-based averaging across ROAs calculated from ROAs within respective cells.

### Expected outcomes

In **Fig. 5** and **Fig. 6**, we illustrate some of the data that STARDUST yields in a classic experimental paradigm. Specifically, acute hippocampal brain slices were obtained from adult mice (P100) in which astrocytes were transduced with AAV5-gfaABC1D::lck- GCaMP6f at P70. GCaMP6f activity in *stratum radiatum* astrocytes was recorded under 2- PLSM. **Fig. 5** serves as a benchmark for STARDUST analysis output from data obtained under different recording conditions: Nikon A1RHD 25 MP microscope (920nm excitation, Nikon 25X, 1.1 NA, 1.6x optical zoom) and Bruker Ultima 2pPlus (920nm excitation, Nikon 25X, 1.1 NA, 1.6x optical zoom). **Fig. 6** provides example data on astrocyte calcium activity in response to the perfusion of low concentrations of norepinephrine (Tocris Biosciences, NE, 200nM). Panels A-B show a classical cell-based averaging of ΔF/F0 and signal properties across ROAs in response to NE with pairwise comparisons. Panels C-H provide examples of ROA-type and ROA-rank based analyses for the experiments shown in A-B.

## Limitations

The current version of STARDUST uses multiple software and platforms, which might prove strenuous for some users or require extra familiarization time. Authors will consider building an all-in-one user-friendly interface in future updates.

## Troubleshooting

**Problem 1:**

In ROA map acquisition (Part 3, step15), large, and connected ROAs arise.

## Potential solution

Reset filter criteria to have a smaller area range in step 12. We encourage users to benchmark the output from a series of area ranges as shown in **Fig. 3** to find the optimal filter criteria.

In addition, users can select a subset of frames to generate ROA maps in step 14 instead of the entire t-stack. This is especially helpful when a long recording was acquired (>15min). We suggest keeping this criterion the same across experiments collected under the same conditions.

## Problem 2

When reading in the cell mask into Jupyter Notebook (step 30), the number of cells does not match the cell number delineated in ImageJ.

## Potential solution

When delineating the cells in step 23, cell boundaries should not touch each other, and the closet points should be a few pixels apart. Adjust cell mask accordingly in ImageJ.

## Problem 3

Recording has a noticeable baseline intensity shift. This issue might become apparent at several steps in the STARDUST pipeline: when visually inspecting the recording, when plotting the z profile in ImageJ (step 3), or during the optional correction step in <u>signal</u> <u>preprocessing</u>section in Part 6 when the slope distribution of all traces is not centered around zero (**Fig. 7A**).

**Figure 7:**
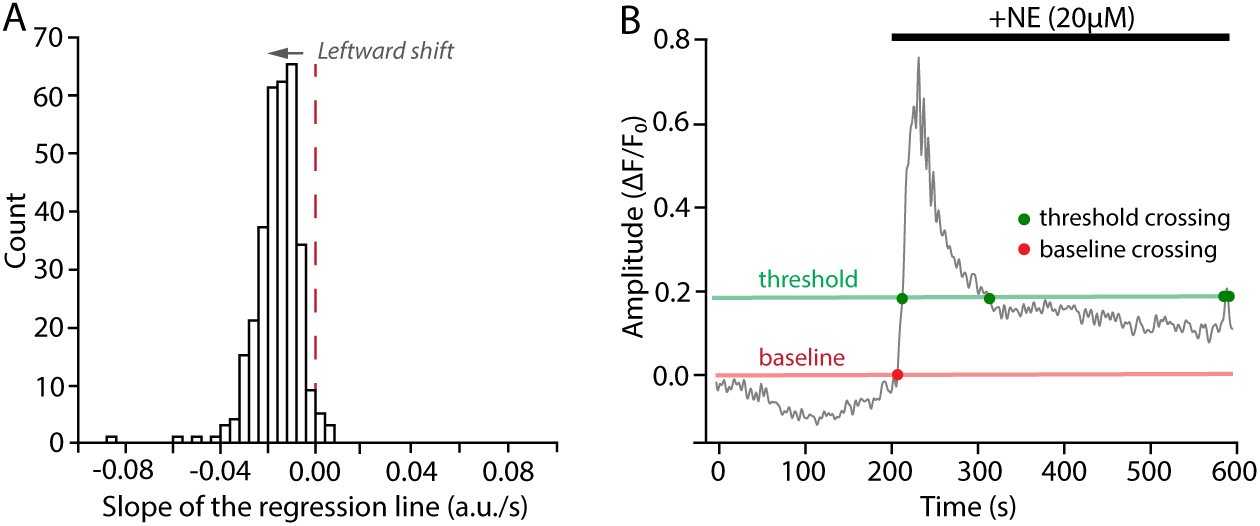
Troubleshooting illustrations. (A) Example of regression slope distribution from a recording with noticeable z-drift. The distribution is centered around -0.015. Also note the lack of Gaussian symmetry (see <u>Troubleshooting 3</u>). (B) Example of a time series in which NE application yielded a sustained signal, with no return to baseline withing the recorded epoch, making baseline fluorescence determination and signal detection difficult (see <u>Troubleshooting 4</u>).

## Potential solution

There are usually possible causes for a baseline intensity shift: 1) physical drifts in x, y, or z axis during the recording, 2) photobleaching, 3) biological change in baseline activities. The first two causes are technical and since both would significantly influence the signal detection, before attempting to compensate for the observed intensity shift, users should carefully examine the recording and decide whether to keep the recording for analysis. If users decide to keep the recording for analysis (in case of minor drifts or photobleaching), apply the optional correction step at signal preprocessing and proceed with the corrected traces for further analysis. Correction factor can be adjusted accordingly from 0 to 1, with 1 being the highest degree of correction. In cases where baseline fluorescence shift might be of biological origin, see case (4) problem below.

## Problem 4

When applying pharmacology, the drug wash-in causes a large response that does not return to F0 before the recording ends, leading to failure in signal detection (since truncated signals are excluded in STARDUST default setting) (see **Fig. 7B**).

## Potential solution

To rescue these truncated signals, users can specify an optional argument *include_incomplete* as True in the ***iterative_baseline*** function (Part 6, Step 33) using the code below. Note that only some of the signal stats, including amplitude, rise time and peak time, will be outputted in the signal analysis summary. Read more about the function and its arguments by using the help documentation.

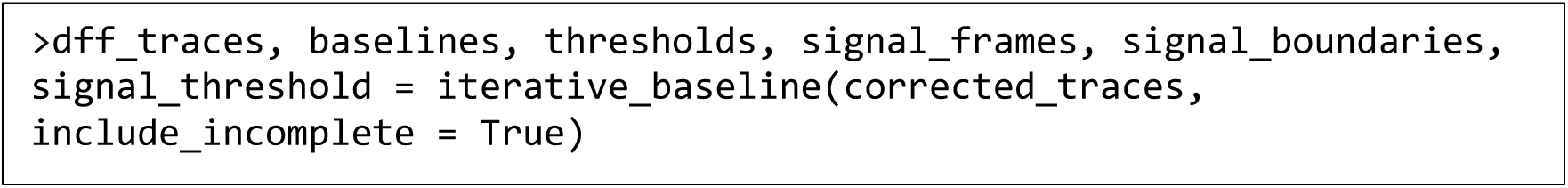

## Problem 5

A large percentage of ROAs are denoted as “inactive” in the output.

## Potential solution

In our conditions, the percent of ‘inactive” ROAs is usually 3-10%. First, make sure the recording is stable and does not have major drifts on x, y, or z axis. If the analyzed recording is stable, a large proportion of “inactive” ROAs might denote 1) suboptimal voxel detection parameters in AQuA, 2) signal threshold being too high, 3) excessive time series with truncated signals (i.e. signals with no initiation or termination points), or 4) significant drifts in the baseline fluorescence, interfering with signal detection.

To identify the cause on a case-by-case basis, visually inspect the trace from these ROAs by using the ***inspect_trace*** function, with a list of ROA IDs, by running the code provided in the Jupiter CheckPoints.

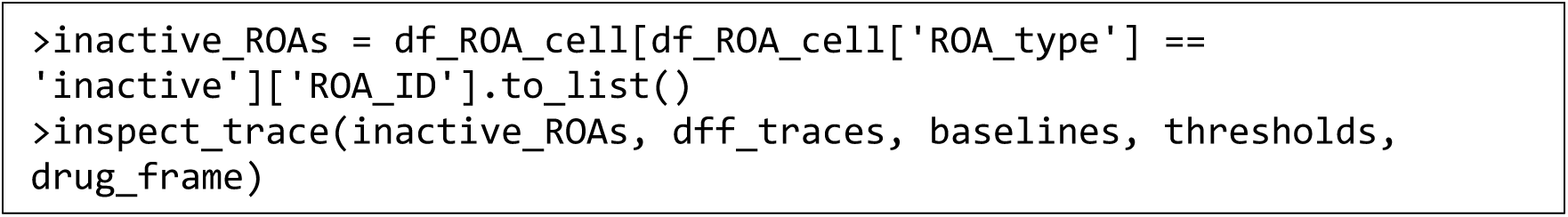

In case (1) where voxel detection parameters are too generous and thus the ROA map included many ROAs without true signals, increase the scaling factor in Part 2 step 10. In case (2) where the signal threshold is too high, rerun the code in the <u>Baseline</u> <u>fluorescence determination and signal detection</u> in Part 6, steps 33-34, and use a lower threshold (for instance, 2SD instead of 3SD) when prompted. In case (3) where the presence of truncated signals is the major cause, refer to Problem 4. Lastly, if the baseline fluorescence shows a noticeable drift (case (4)), consider applying a signal correction at the signal preprocessing step or other smoothening method such as sliding window for F0 determination (not currently supported by the pipeline).

## Article info Resource availability Lead contact

Further information and requests for resources should be directed to and will be fulfilled by the lead contact, Thomas Papouin.

### Technical contact

Questions about the technical specifics of performing the protocol should be directed to and will be answered by the technical contact, Yifan Wu.

### Materials availability

This study did not generate new unique reagents.

### Data and code availability

All original code has been deposited at https://github.com/papouinlab/STARDUST.

## Acknowledgments

T.P. was supported by the National Institutes of Health (1R01MH127163-01), the DoD (W911NF-21-1-0312), the Brain & Behavior Research Foundation (NARSAD Young Investigator Award 28616), the Whitehall Foundation (2020-08-35) and the McDonnell Center for Cellular and Molecular Neurobiology Award (22-3930-26275U). The astrocyte 2-PLSM recordings were performed at the Washington University Center for Cellular Imaging (WUCCI) supported by Washington University School of Medicine, The Children’s Discovery Institute of Washington University and St. Louis Children’s Hospital (CDI-CORE-2015-505 and CDI-CORE-2019-813) and the Foundation for Barnes-Jewish Hospital (3770 and 4642). Graphical abstract was created with <u>BioRender.com</u>.

## Author contributions

Y.W. led the project, developed the pipeline, analyzed data, wrote the first manuscript draft and made the figures. Y.D. developed the interactive version of the pipeline, revised the code and edited the manuscript. K.B.L. contributed example data and provided feedback for the pipeline. T.H. provided feedback for the pipeline. T.P. supervised the project, edited the manuscript and made the figures.

